# UniOP: a universal operon prediction for high-throughput prokaryotic (meta-)genomic data using intergenic distance

**DOI:** 10.1101/2024.11.11.623000

**Authors:** Hong Su, Ruoshi Zhang, Johannes Söding

## Abstract

The study of the deluge of metagenomic and genomic sequences is challenging due to the severe lack of function information. Predicting operons, groups of functionally related genes in prokaryotic genomes, is critical for bridging this gap. However, existing methods for operon prediction heavily rely on experimental data, functional annotations, or extensive characterization of homologous genes, making it difficult to accurately predict operons in newly sequenced or poorly characterized genomes. Here, we introduce UniOP, an unsupervised approach that uses a statistical model to predict operons from intergenic distances directly derived from the target genomic sequence. UniOP not only outperforms alternative approaches on ten complete genomes but also shows superior results on 3269 metagenome-assembled genomes across 13 bacterial and 2 archaeal phyla. Furthermore, we explored enhancing UniOP by incorporating the conservation of gene neighborhood and strandedness in respective genomes and examined the influence of Pfam annotations and motif searching on its performance.

## Introduction

The progress in next-generation sequencing has accelerated the generation of extensive genomic and metagenomic data sets, but the dearth of functional information constrains the analysis of metagenomic data [1]. Even in well-studied ecosystems like the human gut, a striking 40% of genes remain without any functional annotation [1, 2]. In other environments, this fraction can drop to 10% [3]. Functionally related genes are often organized into operons in prokaryotes and in some eukaryotes, such as fungi and nematodes [4, 5]. An operon is a functional unit of DNA consisting of one or more genes transcribed together as a single mRNA molecule under the control of a single promoter [6, 7]. It is estimated that over half of the protein-coding genes in typical bacterial genomes are part of multi-gene operons [8]. Operons play an important role in regulating the expression of genes involved in the same biological process and help prokaryotes adapt to changing environmental conditions. Predicting operons aids in genome function annotation, particularly for newly sequenced genomes where a significant portion of genes may have unknown functions.

Most methods for operon prediction rely heavily on experimental information or functional information, such as microarray gene expression data [9], RNA-seq data [10, 11], common functional annotation [12], or functional associations from STRING scores [13]. These approaches commonly employ probabilistic models [9, 10, 12], or train models based on well-studied *Escherichia coli* and *Bacillus subtilis* organisms with experimentally verified operons [13, 11]. However, they are less reliable when applied to non-model organisms or newly sequenced genomes lacking such information.

Alternatively, some methods identify operons by incorporating the features extracted from genomic sequences, such as intergenic distances, with prior knowledge of operons, including conserved gene clusters, gene order conservation, clusters of orthologous groups, and the frequency of a specific DNA motif in the intergenic region [14, 15, 16, 17]. Many tens of thousands of genomes and metagenome-assembled genomes have become available recently [18], therefore, the methods based on the conservation of gene neighborhoods are getting more powerful. However, they are still biased and limited by the quality, diversity, and number of related genomes.

Deep learning techniques have also been applied to predict operons, including tools like Operon-mapper [11], Operon Hunter [19] and Operon Finder [20]. Operonmapper, based on an artificial neural network model, was trained on verified operons in *Escherichia coli* and *Bacillus subtilis* and uses both intergenic distances and a functional relationship score as input [13]. Operon Hunter predicts operons by utilizing PATRIC[21] to convert genomic data into visual representations, aligning the query genome with reference genomes. These images are then fed into a neural network based on the ResNet18 model, which analyzes gene relationships to make operon predictions. Operon Finder improves on this by using a more efficient MobileNetV2 model, which speeds up predictions and makes the tool accessible on CPUs while maintaining accuracy. Both Operon Hunter and Operon-mapper demonstrate comparable performance, but Operon-mapper slightly outperforms Operon-Hunter in sensitivity [19]. Nevertheless, these methods rely on experimentally verified operons and functional associations from the STRING database [22, 23] for training, which limits their applicability to organisms with extensive functional data.

Due to the scarcity of experimental data, functional information, or related genomes for metagenomic contigs and metagenome-assembled genomes, most wholegenome operon prediction methods are not applicable to metagenomic data. To our knowledge, MetaRon [24] is the first computational pipeline specifically designed for metagenomic data without any experimental or functional information, identifying operons based on co-directionality, intergenic distance, and promoter presence/absence. However, this downloadable software is currently no longer functional.

Here, we present UniOP, an unsupervised probabilistic approach for operon prediction using only intergenic distances derived from the genomic sequence of interest. UniOP is highly efficient, averaging 1.3 seconds per genome on CPU, relying solely on the genomic sequence. Testing on genomes and metagenome-assembled genomes (MAG) indicates that UniOP achieves competitive performance with state-of-the-art methods, offering a robust, scalable solution for operon prediction, particularly in newly sequenced or metagenomic data where traditional approaches fall short.

## Results

### Overview of UniOP

The main steps of the UniOP algorithm are outlined in Fig. 1, with a detailed description in the Methods section. UniOP takes as input a genomic sequence in a FASTA file, containing either a prokaryotic genome or a MAG (Fig. 1A). The Prodigal v2.6.3 program [25] is used to predict the protein-coding sequences from the input genome. Alternatively, users can provide custom protein-coding sequences in Prodigal format.

**Fig. 1.**
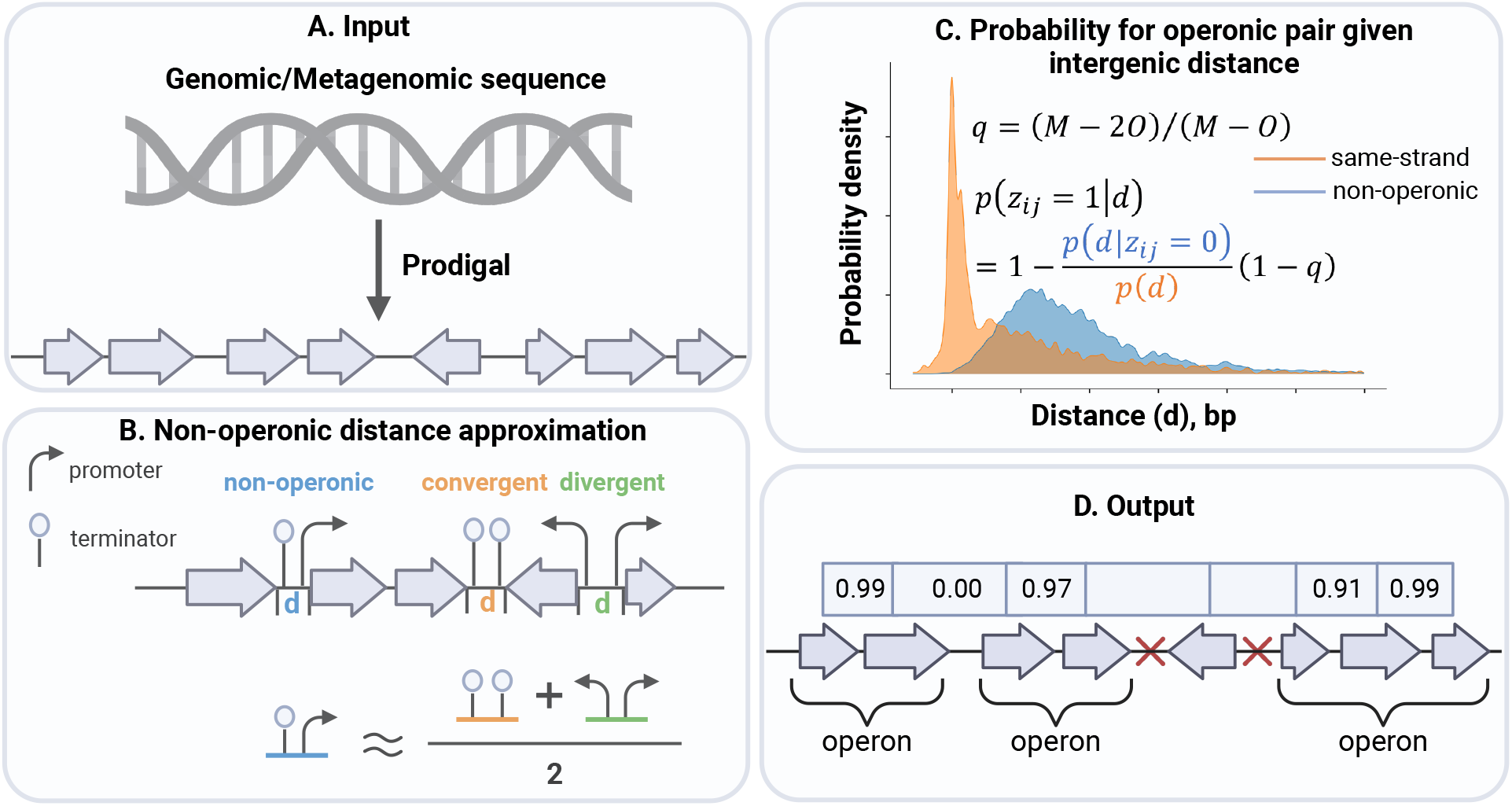
UniOP algorithm. **A**. Predicting protein-coding genes from the input data. **B**. Generating approximated intergenic distances for non-operonic pairs. **C**. Probability for operonic pair given intergenic distance. *p*(*d*) and *p*(*d* | *z*_*ij*_ = 0) are the distance distribution for all same-strand and non-operonic gene pairs, respectively. *q* is a prior probability, estimated by the total number of adjacent same-strand pairs *M* and the number of opposite-strand pairs *O*. **D**. Reporting predictions for whether each same-strand adjacent pair belongs to the same operon. Red fork symbols indicate neighboring genes are on opposite strands, which are excluded from operon analysis.

Our model consists of three main components. First, we approximate a representative list of non-operonic intergenic distances (Fig. 1B). The intergenic space between inter-operon genes typically contains one terminator and one promoter. In contrast, the distance between divergent pairs (neighboring genes on opposite DNA strands transcribed away from each other) contains two promoters, while the distance between convergent pairs (neighboring genes on opposite DNA strands transcribed towards each other) contains two terminators. To approximate non-operonic intergenic distances, we sample 10^4^ random pairs of distances with one distance from the convergent and one from the divergent list, and we take their arithmetic means.

Second, we estimate the prior probability *q* = *p*(*z*_*ij*_ = 1) that two neighboring same-strand genes with indices *i* and *j* = *i* + 1 are part of the same operon. An estimate for *q* can be derived from the number of *M* adjacent gene pairs across all contigs and the number *O* of gene pairs on opposite strands, *q* = (*M* − 2*O*)*/*(*M* − *O*) (Methods, (Fig. 1C)).

Third, we apply an empirical distribution function and kernel density estimation with a Gaussian kernel to estimate the distance distribution for non-operonic pairs and all neighboring gene pairs on the same strand (Fig. 1C). Using Bayes’ theorem, we compute the probability that two neighboring genes belong to the same operon. The algorithm outputs the probabilities that adjacent genes on the same strand belong to the same operon, as well as the predicted operons (Fig. 1D).

### Benchmark UniOP on reference genomes

We assessed the efficiency and accuracy of UniOP on ten genomes with extensively annotated operons (see Table 1). These organisms represent a diverse range of taxa and vary in the number of annotated proteins, as well as in the counts of same-strand adjacent gene pairs, operonic gene pairs, and non-operonic gene pairs. UniOP was applied to either the complete genome or individual chromosomes of each organism, obtained from the NCBI GenBank FTP site [26]. For every genome in the reference dataset, we identified all pairs of genes within annotated operons. We employed an approach utilized in previous operon prediction studies to identify non-operonic gene pairs [13]. Gene pairs (*i, i* + 1)were classified as non-operonic if *i* was part of an annotated operon but *i* + 1 on the same strand was not included in the operon or if *i* + 1 was part of an annotated operon but *i* on the same strand was not (Fig. S7). This definition of non-operonic pairs is preferred over considering all adjacent gene pairs on the same strand outside of operons as non-operonic. The reason is that this approach reduces the occurrence of false negatives, especially when operon annotations are incomplete where genes that are supposed to be part of an operon but not fully annotated.

**Table 1.** Species used to benchmark UniOP.

| Species | Phylum | #annotated proteins | #adjacent pairs | #operonic pairs | #non-operonic pairs |
| --- | --- | --- | --- | --- | --- |
| *Escherichia coli* | Proteobacteria | 4319 | 3028 | 1757 | 671 |
| *Sinorhizobium meliloti* | Proteobacteria | 3392 | 2309 | 969 | 580 |
| *Campylobacter jejuni* | Proteobacteria | 1877 | 1499 | 1028 | 303 |
| *Salmonella enterica* | Proteobacteria | 4537 | 3208 | 1595 | 775 |
| *Bacillus subtilis* | Firmicutes | 4226 | 3080 | 1836 | 724 |
| *Clostridium beijerinckii* | Firmicutes | 5252 | 4108 | 1174 | 1048 |
| *Clostridioides difficile* | Firmicutes | 3819 | 3131 | 1296 | 900 |
| *Mycobacterium tuberculosis* | Actinobacteria | 4085 | 2695 | 1364 | 650 |
| *Eggerthella lenta* | Actinobacteria | 3116 | 2276 | 1067 | 641 |
| *Synechococcus elongatus* | Cyanobacteria | 2670 | 1724 | 707 | 402 |

### Estimated *q* values are reliable

The prior probability that adjacent genes on the same strand belong to an operon, *q* is an important parameter of the model that needs to be accurately estimated. We compared annotated and estimated values of *q* across the ten genomes to evaluate the accuracy. The annotated value of *q*_annot_ is calculated by dividing the number of annotated operonic gene pairs by the total count of adjacent gene pairs on the same strand. Conversely, the estimated value of *q*_est_ is determined by dividing the estimated number of operonic gene pairs, as outlined in the Methods section, by the total number of adjacent gene pairs on the same strand. Fig. 2A and Table S1 present the comparison results for the relative deviation, log(*q*_est_*/q*_annot_). Notably, for the well-studied genome *Escherichia coli*, which has the most extensively annotated operons, the relative error (ratio in Table S1) between the estimated and annotated *q* values is -0.011, indicating the estimated value is approximately 97.7% of the annotated value. Other well-studied genomes such as *Mycobacterium tuberculosis* and *Bacillus subtilis* exhibit similar accuracy, supported by the minimal error bars in Fig. 2A.

**Fig. 2.**
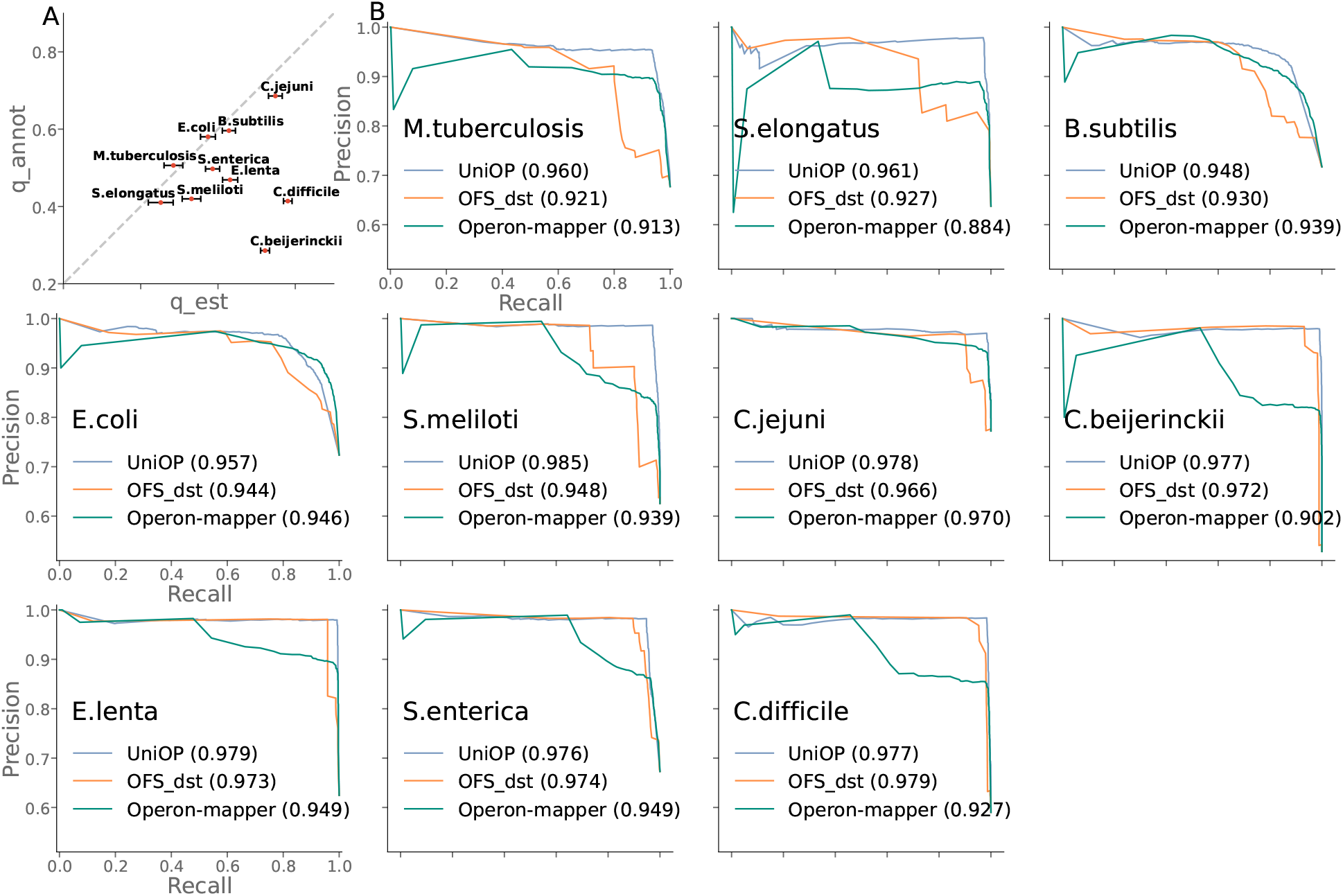
The prediction performance on annotated reference genomes. **A**. Comparison between annotated and estimated *q* values. The horizontal lines with caps on each point are the error bars representing the uncertainty in estimated *q*. **B**. Precision-recall curves comparing the performance of UniOP, OFS dst, and Operon-mapper. Operon-mapper predictions are fetched from its web server, while OFS dst predictions are locally acquired by running its script. Values in parentheses are AUC-PR. All subplots have the same x-axis and y-axis scales.

Based on the relative deviation of *q* values, the ten examined genomes are categorized into three broad groups, as shown in Table S1. The first group includes genomes *Escherichia coli, Mycobacterium tuberculosis, Bacillus subtilis, Campylobacter jejuni*, and *Synechoccus elongatus*, with minimal relative deviations (−0.011 to 0.097), indicating complete operon annotations. The second group, comprising *Salmonella enterica, Sinorhizobium meliloti*, and *Eggerthella lenta*, shows larger deviations (0.164 to 0.298), suggesting relatively incomplete annotations. The last group, *Clostridioides difficile* and *Clostridium beijerinckii*, with significant deviations (0.643 to 0.926), implies substantial gaps in operon annotations. This increases the likelihood of misclassifying operonic pairs as non-operonic pairs as incomplete operon annotations can result in one gene from a supposed operonic pair being annotated while the other is not (see Methods section and Fig. S6).

### UniOP shows homogenous performance across a diverse set of prokaryotic genomes

OFS and Operonmapper are two available methods for operon prediction [12, 11]. OFS, an unsupervised method, predicts operons by integrating intergenic distance, conserved gene clusters, and functional relatedness. It exhibits promise for operon prediction in genomes with limited annotation, as it relies on fewer data sources than other approaches. However, the **cluster finder.pl** script, which identifies the conserved gene clusters, is unavailable due to dependencies on multiple outdated external programs. Furthermore, not all microbial genomes, especially those newly sequenced, possess function annotations. Hence, we compared UniOP with the distance model of OFS, denoted as OFS dst. UniOP and OFS dst both assess the likelihood of adjacent gene pairs belonging to the same operon by applying Bayes’ theorem (see Methods section). They differ in their strategies for generating the non-operonic pairs and estimating *q* (the prior probability that two adjacent genes on the same strand are in a common operon), *p*(*d*) (the length distribution of intergenic distances within the same strands), and *p*(*d* | *z*_*ij*_ = 0) (the length distribution of intergenic distances when adjacent genes on the same strand are not part of the same operon). Operon-mapper, a recent machine learning-based method, achieves similar prediction accuracy to Operon Hunter and Operon Finder [19], with a more user-friendly interface. Consequently, we compared UniOP against OFS dst and Operon-mapper.

Fig. 2B presents precision-recall curves of UniOP, OFS dst, and Operon-mapper across ten genomes, grouped into three categories based on the relative deviation in *q*. UniOP consistently demonstrates high precision and recall across diverse prokaryotic genomes, underscoring both accuracy and broad applicability. Notably, UniOP performs well even in the third group, where substantial gaps in operon annotations lead to the false classification of operonic pairs as non-operonic. This robustness is due to the stringent definition of non-operonic pairs, which effectively reduces falsely annotated negatives.

Across the ten genomes, UniOP achieves higher AUC-PR scores than OFS dst in nine cases and significantly outperforms Operon-mapper on all genomes, including *Escherichia coli* and *Bacillus subtilis*, for which Operon-mapper was specifically trained. In these two cases, OFS dst fails to surpass Operon-mapper, further highlighting UniOP’s improved and robust performance. Un-like OFS dst and Operon-mapper, which show sharp declines in precision as recall increases, UniOP maintains consistently high precision even at high recall values, indicating superior discrimination between operonic and non-operonic gene pairs.

UniOP performs exceptionally well on genomes in the first group, such as *Mycobacterium tuberculosis, Syne-choccus elongatus, Bacillus subtilis, Escherichia coli*, and *Campylobacter jejuni*. For the genomes in the second and third groups, UniOP’s AUC-PR values are only marginally higher than OFS dst, with a slight 0.002 lower AUC-PR on *Clostridioides difficile*. This is likely due to two factors: (i) misclassifications within the ground truth set, where even stringent criteria for non-operonic pairs cannot fully eliminate classification errors, and (ii) OFS dst’s limited ability to distinguish between positives and negatives, resulting in more incorrect predictions. Factor (i) may artificially inflate OFS dst’s evaluation scores. The evidence of misclassification of operonic pairs as non-operonic in third-group genomes is further reflected by UniOP’s lower precision and specificity on *Clostridioides difficile* (light purple bar in Fig. S1B) and *Clostridium beijerinckii* (dark red bar in Fig. S1B), as opposed to its higher scores on other genomes, particularly those in the first group.

### Impact of conserved gene clusters

Genes are often functionally associated if clusters of homologous genes with similar arrangements are found in many genomes [14, 27]. The consistent pattern across multiple genomes (denoted as conserved gene clusters) suggests that evolutionary pressure keeps these genes together, indicating that they probably participate in the same function. To examine the impact of conserved gene clusters on our approach, we trained a logistic regression model for each query genome, using the conservation of gene neighborhood and strandedness in related genomes as input features. Details on conservation information can be found in the Methods section and Supplementary materials.

The training samples consist of gene pairs with confident predictions from UniOP. We classify the top *αq* fraction of gene pairs with the highest probabilities as operonic pairs and the bottom *α*(1 − *q*) fraction with the lowest probabilities as non-operonic. The parameter of *α* controls the proportion of gene pairs that receive a label, with a smaller *α* yielding fewer but more reliable pseudo labels. In this study, we set *α* = 0.5.

We trained two logistic regression models using two conservation matrices: the raw conservation matrix, derived from the conserved gene clusters in reference genomes, and the refined matrix, generated by applying UniOP to each reference genome and updating the raw matrix with UniOP predictions. The final prediction combines UniOP predictions with the logistic regression models by including them as a bias term, using fixed weights trained on the conservation matrices.

The results in Fig. S2 reveal that integrating intergenic distance with the raw conservation matrix enhances performance in six out of ten evaluated genomes (Comparing UniOP to UniOP cons raw). Moreover, combining the intergenic distance model with the refined conservation matrix further improves performance in eight out of ten genomes (UniOP cons raw versus UniOP cons). However, as depicted in Fig. S3, the inclusion of conservation data slows down the tool significantly, making large-scale applications more challenging. We have therefore decided not to incorporate conserved gene cluster information in our final approach.

### Comparative evaluation of runtime

Runtime is essential in assessing a tool’s suitability for rapidly annotating a vast collection of genomes in a highthroughput manner. Given that the Operon-mapper’s predictions depend on its web server, the total running time includes queuing time, causing a single genome to take several hours. This introduces inefficiencies when analyzing a large set of genomes. We therefore focus on the runtime comparison between UniOP and OFS dst on the ten genomes, as depicted in Fig. S3. The measurements were made using the same hardware: 8-core, 16-thread Intel 11th Gen Core i7-11700K @3.6 GHz (5.0 GHz max boost).

Fig. S3 shows that OFS dst consistently exhibits fast and stable processing speeds, completing predictions in approximately 0.24 seconds on average. In contrast, UniOP operates at a slower pace, with runtime scaling linearly with the number of genes within a genome. For instance, *Clostridium beijerinckii* has nearly 2.8 times more genes than *Campylobacter jejuni*, and UniOP’s runtime increases proportionally (1.99 seconds vs 0.71 seconds), reflecting a less efficiency with larger genomic datasets compared to OFS dst. Nonetheless, UniOP’s overall runtime remains competitive, with an average prediction time of 1.3 seconds.

### Application to human gut MAGs

To examine the applicability of UniOP on metagenomic data, we applied it to 3269 non-redundant and diverse human gut MAGs from 15 prokaryotic phyla (13 bacterial and 2 archaeal). These genomes were downloaded from human gut microbiota, with many of the species being uncultured [2]. The analysis by UniOP revealed a total of 1,603,492 operons, encompassing 5,717,497 genes across all genomes. On average, about 68.7% of genes within each phylum were predicted to be part of operon structure. For detailed information on annotations and predictions for each phylum, refer to Table S2.

In the absence of experimental operon annotation for all MAGs, directly evaluating the accuracy of predicted operons is challenging. Nonetheless, an indirect assessment can be conducted using KEGG module annotations [28] from the same study that provided the MAGs [2]. To evaluate the performance of predicted operons within each MAG, we focus on pairs of neighboring genes for which both genes have at least one KEGG module annotation. The column labeled **%annotated pairs** in Table S2 illustrates the proportion of gene pairs within each phylum annotated by the KEGG modules, with values ranging from 0.082 to 0.153. This reveals a notable scarcity of gene pairs annotated by these modules, highlighting the significant challenge posed by the severe lack of functional annotation in metagenomic analysis.

### Evidence supporting the reliability of predicted operons by UniOP in MAGs

Fig. 3 presents the predictive performance of UniOP and OFS dst across all annotated gene pairs within 15 phyla. Gene pairs sharing at least one common KEGG module are annotated as operonic pairs (positives), while those from different KEGG modules are labeled as non-operonic (negatives). The figure reveals that UniOP performs well on 10 out of the 15 phyla, achieving AUC-PR scores between 0.81 to 0.94, and outperforms OFS dst in 11 out of the 15 phyla. At the threshold of 0.5, UniOP also demonstrates higher precision and specificity than OFS dst across all phyla (Fig. 3A, Fig. S4), highlighting its effectiveness in distinguishing operonic from non-operonic pairs.

**Fig. 3.**
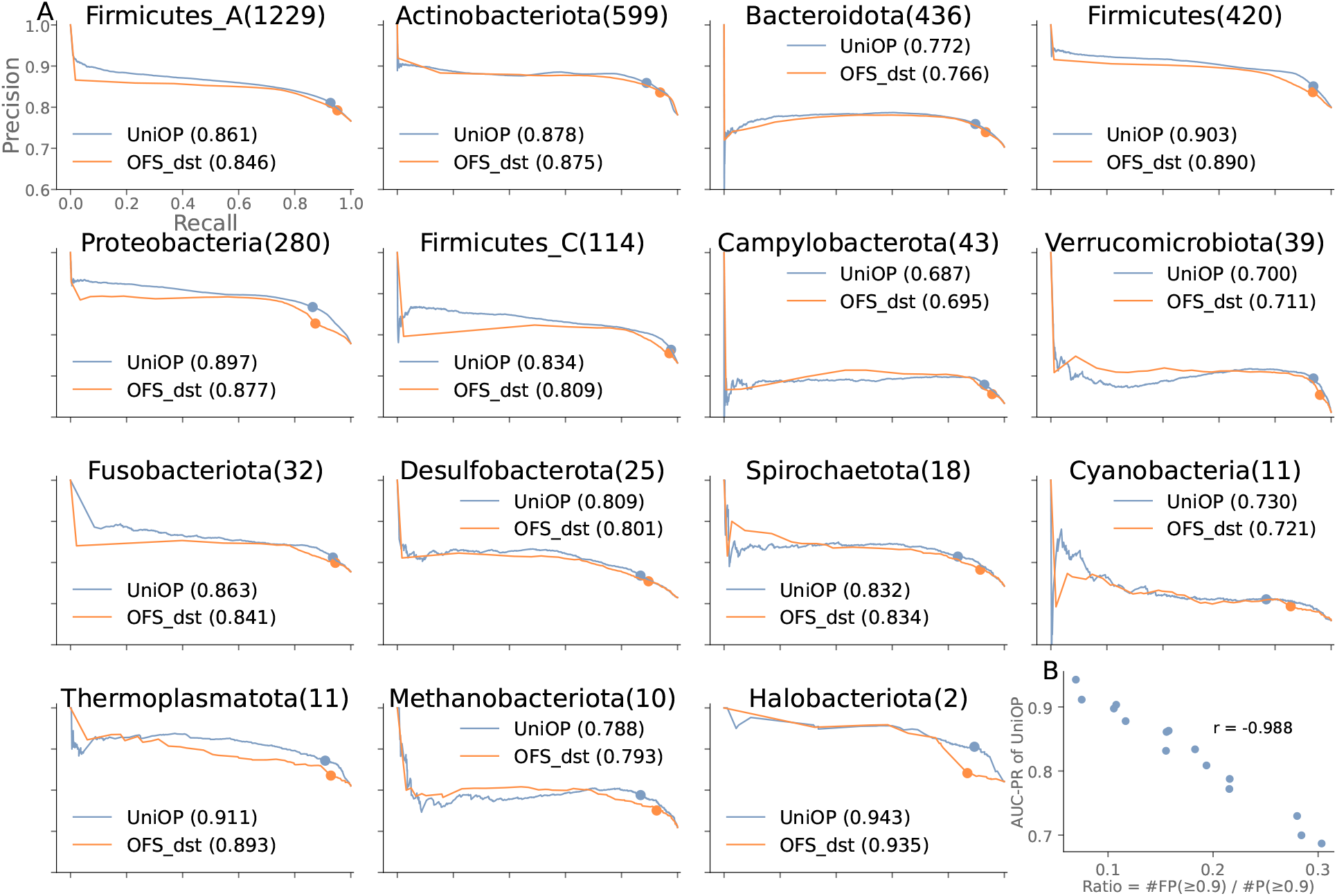
The predictive performance on MAGs. **A**. Comparison of Precision-Recall curves of UniOP and OFS dst. There are 13 bacterial and 2 archaeal phyla (the last two subplots). Each dot on the curve indicates the precision and recall values at a threshold of 0.5. The number in the title indicates the total number of MAGs in each phylum. The legend shows the AUC-PR scores for each method. The x-axis and y-axis ranges are consistent across all subplots. **B**. Correlation between the proportion of predictions with probability ≥ 0.9 that are false positives (FPs) and AUC-PR for UniOP.

Upon examining the negative pairs falsely predicted as positives (FPs) of UniOP for all phyla, we observed that AUC-PR scores across 15 phyla highly correlate (Pearson r=-0.988) with the ratio of FPs with a probability ≥ 0.9 to the predictions with a probability ≥ 0.9, as illustrated in Fig. 3B. The five phyla with the highest FP ratios have the lowest AUC-PR values (0.69 to 0.79). This may result from functionally associated genes that are not labeled with the same KEGG pathway ID, which leads to a higher rate of falsely annotated operonic pairs as non-operonic, thus underestimating the predictive performance of UniOP in these cases.

### Conservation analysis of MAGs operon

To gain more insights into the predicted operons for MAGs, we focused on the phylum Firmicutes A as a case study. Firmicutes A, the largest phylogenetic group, includes 1229 MAGs from uncultured species. We started by assembling the predicted operonic pairs into complete operons for analysis.

For the computation of gene clusters we used MMseqs2 with thresholds: minimum sequence identity 50%, *E* - value cutoff 0.001, and minimum sequence coverage 80%. The parameters used were `--min-seq-id` 0.5 -c 0.8 `--cov-mode` 0 -e 0.001. Conservation scores for each operon were calculated based on these clusters and categorized into four distinct types. For clarity, consider two operons, *A* and *B*, where the conservation type of operon *A* to operon *B* is as follows.

- **Perfect Match**: When the genes within operon *A* exhibit a one-to-one homologous relationship with the genes in operon *B*, and both operons are of equal size, *B* is considered a ‘perfect’ match to *A*.
- **Under Match**: If operon *A* has fewer genes than operon *B*, yet each protein in operon *A* possesses a corresponding homolog in operon *B*, this scenario is termed as an ‘under’ match.
- **Over Match**: Conversely, if operon *A* is larger than operon *B*, and each protein in operon *B* has a corresponding homologous in operon *A*, this condition is labeled as an ‘over’ match.
- **No Match**: In all other instances, we assign the ‘No’ match category for operon *A*.

It’s important to note that an operon from one MAG may align with multiple operons in another MAG. In such cases, ‘perfect’ matches always take priority, followed by ‘under’ matches, and then ‘over’ matches. Based on these definitions, we calculated the fraction of each type. Overall, approximately 46.7% of operons exhibit one or more ‘perfect’ matches in other MAGs, about 8.3% belong to the under-match group, and an additional 15.7% belong to the over-match group. Fig. S5A shows the distribution of conserved operons in each category according to the length of the operons. The number on the top of each bar represents the total number of operons with a specific length. As illustrated in the figure, as the length of the operons increases, the fraction of operons with ‘perfect’ and ‘no’ matches decreases, while the fraction of operons with ‘over’ matches increases. The observed trends may be attributed to the complexity of longer operons, which are prone to extensive evolutionary changes such as gene loss, horizontal gene transfer, or rearrangements, all of which can disrupt their conservation.

To ascertain whether these observations are statistically significant or merely chance occurrences, we established a baseline for comparison. This was achieved by substituting the genes in each operonic pair with two randomly selected non-operonic genes. The resulting recalculated conservation outcomes, as depicted in Fig. S5B, exhibit a markedly different distribution compared to the original dataset. This notable contrast implies that the observed patterns in the actual data likely stem from underlying biological processes and evolutionary pressures acting on the operons.

## Discussions

We developed UniOP, an unsupervised statistical model for accurately inferring operons in prokaryotic genomic and metagenomic data. UniOP relies solely on intergenic distance between neighboring genes, making it universally applicable to large-scale datasets.

Our evaluation demonstrated UniOP’s high accuracy and efficiency across ten diverse, well-annotated prokaryotic genomes, achieving AUC-PR values ranging from 0.948 to 0.985. UniOP reliably distinguishes between operonic and non-operonic gene pairs by assigning high probabilities to operonic pairs and low probabilities to non-operonic ones, maintaining stable proportions regardless of the binary classification threshold. Decently high AUC-PR scores of UniOP across 15 non-redundant prokaryotic phyla (13 bacterial and 2 archaeal) despite patchy KEGG annotation further demonstrates its generalizability to metagenomes-assembled genomic sequences.

We explored the influence of conserved gene pairs on UniOP’s performance. While this information enhanced prediction accuracy, the time-intensive nature of identifying conserved gene pairs limits the scalability of largescale applications.

We also investigated the potential benefit of incorporating DNA motif information, considering that certain DNA motifs seem to be often associated with inter-operonic regions [16]. We approached this by (i) training a convolutional neural network on intergenic regions to predict promoters, where DNA motifs are typically found, and (ii) detecting motifs using Bammmotif [30, 31]. We then incorporated the predicted promoters or motifs into the logistic regression model (see the “Impact of conserved gene clusters” section) as additional features. However, neither approach significantly improved UniOP’s performance.

Additionally, we assessed the integration of Pfam domain annotations, given that certain domains often cooccur within the same operons due to shared molecular functions. Using **hmmscan** http://hmmer.org/, we searched against the Pfam profile database [32]. The resulting Pfam annotations were incorporated into UniOP by multiplying the Bayes factors [33], but this did not yield further improvements.

UniOP stands out from state-of-the-art methods by achieving remarkable accuracy using only the intergenic distances, eliminating the need for experimental or functional annotations. This ensures universal applicability across all prokaryotic genomes, including those assembled from metagenomic data. It enables rapid processing of large-scale datasets, making it suitable for extensive studies. As user-friendly command-line software compatible with Linux and macOS, UniOP requires only a genomic sequence file in FASTA format or a proteincoding gene annotation file in GFF3 format. Its minimal external dependencies ensure stability and simplify installation and maintenance, empowering researchers to unlock insights with unprecedented speed and precision.

## Materials and Methods

### Same-operon probability given the intergenic distance

The intergenic distance, defined as the number of base pairs between two adjacent genes, is important in predicting operons. Genes within the same operon typically exhibit a smaller intergenic distance than genes not belonging to the same operon [34, 35]. This data source is available to all prokaryotic (meta-)genomes. In this study, our objective is to determine the likelihood that two consecutive same-strand proteins are part of the same operon, given their intergenic distance. Applying Bayes’ theorem, this probability can be expressed as:

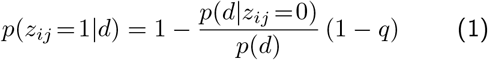

Here, *i* and *j* indicate gene indices, where *j* = *i* + 1 and two genes are on the same strand. The intergenic distance *d* is calculated as *j* start − *i* stop + 1, where *j* start and *i* stop denote the start position of gene *j* and the stop position of gene *i* on the genomic sequence, respectively. The indicator variable *z*_*ij*_ takes the value 1 if genes *i* and *j* belong to the same operon and 0 otherwise. The variable *q* = *p*(*z*_*ij*_ = 1) represents the global prior probability that two neighboring same-strand genes are part of the same operon.

By accurately estimating three key probabilities *q, p*(*z*_*ij*_ = 0|*d*), and *p*(*d*), we can determine the likelihood that two consecutive same-strand genes belong to the same operon based on their intergenic distance, *p*(*z*_*ij*_ = 1|*d*).

### Estimation of the prior probability *q*

To estimate *q*, the fraction of consecutive genes on the same strand belonging to the same operon, consider a genome consisting of one or more contigs. Each contig contains a collection of annotated genes, where each contig’s first and last genes may be partial. We introduce variables to facilitate the calculation. Let *S* denote the number of same-strand adjacent gene pairs across all contigs, and *O* represent the number of opposite-strand adjacent gene pairs. The total number of adjacent gene pairs across all contigs is denoted as *M* = *S* + *O*. The same-strand adjacent gene pairs *S* can be further decomposed into intra-operonic pairs *I* and inter-operonic pairs *R*, such that *S* = *I* + *R*. We can estimate the expected number of inter-operonic gene pairs as approximately equal to the expectation of opposite-strand pairs (i.e., *R* ≈ *O*) under the assumption that genes are randomly distributed and the orientation of adjacent genes on opposite strands is independent of operon structure. Finally, the fraction *q* is calculated as the ratio of the number of intra-operonic gene pairs (*I*) to the total number of the same-strand adjacent gene pairs (*S*). Hence, we estimate the prior probability as follows:

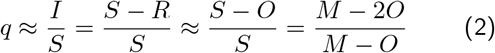

### Estimation of *p*(*d*) and *p*(*d*|*z*_*ij*_ = 0)

#### Generation of non-operonic intergenic distances

To estimate the distribution *p*(*d* | *z*_*ij*_ = 0), where *z*_*ij*_ = 0 represents the adjacent gene pairs belonging to different operons, we need to compute intergenic distances for these inter-operon pairs. Given the frequent lack of operon annotations, we developed a method to approximate these distances applicable across all prokaryotes. Typically, the intergenic region between inter-operon genes includes one terminator and one promoter [36]. In contrast, the distance between divergently transcribed genes (neighboring gene pairs on opposite DNA strands transcribed away from each other) contains two promoters, while the distance between convergently transcribed genes (neighboring gene pairs on opposite DNA strands transcribed towards each other) contains two terminators.

Based on these observations, we generated a representative list of non-operonic intergenic distances by randomly sampling pairs of distance, one from the convergent list and one from the divergent list, and calculating the arithmetic mean of each pair. To ensure the robustness of our approach, we sample 10^4^ pairs per genome, resulting in a curated list that encompasses diverse combinations of convergent and divergent intergenic distances. Each distance in this list represents the average length of one promoter and one terminator.

This curated list allows us to approximate the distribution *p*(*d*|*z*_*ij*_ = 0) in a way that captures the intricacies of inter-operon intergenic distances.

#### Approximation of *p*(*d*) and *p*(*d*|*z*_*ij*_ = 0)

To obtain reliable probability densities from the empirical values, we employ the kernel density estimation approach with a Gaussian kernel. However, for large distances, both densities of *p*(*d*) and *p*(*d* | *z*_*ij*_ = 0) become extremely low, resulting in significant errors in the calculation of 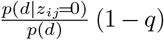. To address this issue, data transformation is performed using an empirical distribution function denoted by *Q*(*d*). This involves ranking all unique observations in the data sample and calculating the *Q*(*d*) as the empirical cumulative probability distribution function for the *d*_*i*_ points:

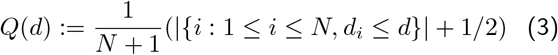

A kernel density estimation with a Gaussian kernel is then applied to the transformed intergenic distances {*Q*(*d*_*i*_)|1 ≤ *i* ≤ *N*} and non-operonic intergenic distances 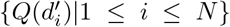, leading to densities 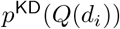 and 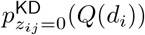. Notably, both variables, *d*_*i*_ and 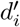, undergo the same monotonous transformation *Q*(*d*). As a result, the factors 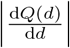 that is multiplied by the density of transformed variable *Q*(*d*) effectively cancel each other out during the transformation process.

Consequently, we have:

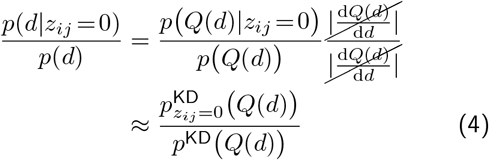

This approximation allows for a more robust calculation, mitigating the challenges of extremely low densities at large distances.

### Generation of conservation matrix

In the absence of selective pressure, prokaryotic genomes tend to undergo rearrangements [37], disrupting neighborhoods. Consequently, genes that cluster together in multiple organisms are more likely to be part of the same operon. To examine this contribution, we compared the predictions of combining the information of conserved gene clusters with UniOP.

Suppose we have a set of reference genomes. For each genome among the ten examined, we searched against the reference genome set to detect gene clusters. Specifically, for each query genome, we first employed the Spacedust method for *de novo* cluster discovery [38] to systematically identify conserved gene clusters across reference genomes. Next, we constructed a raw conservation matrix, where each row represents a genome in the reference set, and each column represents a samestrand neighboring gene pair in the query genome. An element in this matrix is set to 1 if the corresponding gene pair in the reference genome is contained in a gene cluster conserved in the genome corresponding to the row. Otherwise, the element is set to 0.

Additionally, we created a refined conservation matrix by substituting values in the raw conservation matrix with the UniOP probabilities for the reference genomes. Specifically, if the element in the raw conservation matrix is 1 and the gene pair in the corresponding genome is adjacent on the same strand, then the value is replaced by the UniOP probability for this operonic pair. Otherwise, the values remain the same. Please refer to the supplementary materials to see more details.

### Evaluation criteria

In this study, we utilized the Area Under the Precision-Recall curves (AUC-PR) as the primary metric to evaluate prediction performance. AUC-PR is generally more informative for imbalanced datasets, where it provides a clearer measure of performance than the more commonly used Area Under Receiver Operating Characteristic curves (AUC-ROC) [39]. Other metrics like Precision = TP*/*(TP+FP), Recall = TP*/*(TP+FN), and Specificity = TN*/*(TN+FP) are also used to assess classifier performance. TP (True Positives) is the number of correctly predicted operonic pairs, FP (False Positives) is the number of non-operonic pairs incorrectly predicted as operonic, TN (True Negatives) represents the number of correctly predicted non-operonic pairs, and FN (False Negatives) denotes the number of operonic pairs incorrectly predicted as non-operonic.

### Error calculation of *q*

To evaluate the precision of *q*, the prior probability given by *q* = (*M* − 2*O*)*/*(*M* − *O*), we calculated the uncertainty of *q* as follows: assuming that *M* and *O* follow Poisson distributions, the uncertainties (standard deviations) are 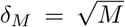 and 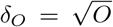 The partial derivatives for error propagation are 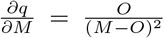 and 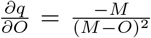 The uncertainty of *q* is then calculated using the propagation of uncertainty formula :

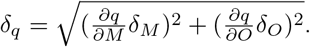

### Benchmark dataset and operon resources

To assess the effectiveness of our operon prediction approach, we conducted tests on ten well-known organisms for their extensively characterized operon data, as experimentally verified operons are not readily available for most microbes. The organisms we examined include *Escherichia coli K-12 MG1655* (GCF 000005845.2), *Bacillus subtilis* (GCF 000009045.1), *Campylobacter jejuni RM1221* (GCF 000011865.1), *Mycobacterium tuberculosis H37rv* (GCF 000195955.2), *Sinorhizobium meliloti 2011* (GCF 000346065.1), *Eggerthella lenta DSM2243* (GCF 000024265.1), *Synechoccus elongatus PCC 7942* (GCF 000012525.1), *Clostridium beijerinckii NCIMB 8052* (GCF 000016965.1), *Clostridioides difficile 630* (GCF 000009205.2), and *Salmonella enterica subsp. enterica Serovar Typhimurium str. 14028S* (GCF 000022165.1). We obtained the complete genome sequences from the NCBI GenBank FTP [26]. Protein sequences and GFF3 format [40] files, which contain information about the locations of annotated proteins, were generated using Prodigal v2.6.3 [25]. For operon annotation, we utilized the RegulonDB database version [41] for *Escherichia coli* and *SubtiWiki* [42] for *Bacillus subtilis. SubtiWiki* was chosen as it provides a considerably larger number of experimentally validated operons than the widely-used DBTBS database [43]. The operon annotation for the remaining eight genomes was obtained from the OperomeDB database [44]. Although the operons in OperomeDB are also predictions, they are characterized by high accuracy and reliability. This high precision is attributed to using available RNA-sequencing datasets, encompassing a diverse array of experimental conditions. For detailed data, please refer to Table 1.

### Metagenomic datasets for comprehensive analysis

We expanded the application of our method to encompass Metagenome-Assembled Genomes (MAGs), showcasing its robustness in handling metagenomic data. The dataset utilized in this analysis was curated from a diverse pool of 4689 species-level human gut prokaryotes. These prokaryotes were initially derived from an extensive collection of 289,232 genomes using a multistep distance-based clustering approach, as detailed in the study by [2]. Subsequently, genomes with contamination *>* 5% or completeness *<* 90% were filtered out, along with phyla containing fewer than 10 MAGs, resulting in a final set of 3269 high-quality MAGs covering 15 phyla. Notably, this set includes two archaeal phyla with ***≤*** 10 high-quality MAGs each, as there were no other archaeal phyla in the original dataset.

### Availability of data and materials

Our approach’s source code, compilation instructions, and a brief user guide are available at https://github.com/hongsua/UniOP.Additionally, the annotated protein files, annotated operon files, extracted operonic gene pairs, and non-operonic gene pairs for genomes in the benchmark dataset are available at https://github.com/hongsua/UniOP/ data/ten_annotated_organisms. KEGG annotations and UniOP predictions for the metagenomic data are also provided in the same repository at https://github.com/hongsua/UniOP/data/mags. The annotated protein files of MAGs used in this study were downloaded from http://ftp.ebi.ac.uk/pub/databases/metagenomics/mgnify_genomes/human-gut/v2.0.2/.

## Acknowledgements

This work used the Scientific Compute Cluster at GWDG, the joint data center of the Max Planck Society for the Advancement of Science (MPG) and the University of Göttingen.

## Contributions

HS and JS designed the UniOP algorithm, benchmark, and biological application. HS and RS developed the algorithm. HS performed benchmarks and generated figures. HS and JS wrote the manuscript. All authors read and approved the final manuscript.

## References

[1] Thomas, A. M. & Segata, N. Multiple levels of the unknown in microbiome research. BMC Biology 17, 48 (2019). URL 10.1186/s12915-019-0667-z.

[2] Almeida, A. et al. A unified catalog of 204,938 reference genomes from the human gut microbiome. Nature biotechnology 39, 105–114 (2021).

[3] Hurwitz, B. L. & Sullivan, M. B. The pacific ocean virome (POV): a marine viral metagenomic dataset and associated protein clusters for quantitative viral ecology. PLoS One 8, e57355 (2013).

[4] Rokas, A., Wisecaver, J. H. & Lind, A. L. The birth, evolution and death of metabolic gene clusters in fungi. Nat Rev Microbiol 16, 731–744 (2018).

[5] Zorio, D. A., Cheng, N. N., Blumenthal, T. & Spieth, J. Operons as a common form of chromosomal organization in c. elegans. Nature 372, 270–272 (1994).

[6] Ermolaeva, M. D., White, O. & Salzberg, S. L. Prediction of operons in microbial genomes. Nucleic acids research 29, 1216–1221 (2001).

[7] Blumenthal, T. Operons in eukaryotes. Briefings in Functional Genomics 3, 199–211 (2004).

[8] Okuda, S. et al. Characterization of relationships between transcriptional units and operon structures in Bacillus subtilis and Escherichia coli. BMC Genomics 8, 48 (2007). URL 10.1186/1471-2164-8-48.

[9] Sabatti, C., Rohlin, L.Oh, M.-K. & Liao, J. C. Co-expression pattern from dna microarray experiments as a tool for operon prediction. Nucleic acids research 30, 2886–2893 (2002).

[10] Tjaden, B. A computational system for identifying operons based on rna-seq data. Methods 176, 62–70 (2020).

[11] Taboada, B., Estrada, K., Ciria, R. & Merino, E. Operonmapper: a web server for precise operon identification in bacterial and archaeal genomes. Bioinformatics 34, 4118– 4120 (2018).

[12] Westover, B. P., Buhler, J. D., Sonnenburg, J. L. & Gordon, J. I. Operon prediction without a training set. Bioinformatics 21, 880–888 (2005).

[13] Taboada, B., Verde, C. & Merino, E. High accuracy operon prediction method based on string database scores. Nucleic acids research 38, e130–e130 (2010).

[14] Overbeek, R., Fonstein, M., D’Souza, M., Pusch, G. & Maltsev, N. The use of gene clusters to infer functional coupling. Proc. Natl. Acad. Sci. USA 96, 2896–2901 (2013).

[15] Che, D., Zhao, J., Cai, L. & Xu, Y. Operon prediction in microbial genomes using decision tree approach. In 2007 IEEE Symposium on Computational Intelligence and Bioinformatics and Computational Biology, 135–142 (IEEE, 2007).

[16] Dam, P., Olman, V., Harris, K., Su, Z. & Xu, Y. Operon prediction using both genome-specific and general genomic information. Nucleic acids research 35, 288–298 (2007).

[17] Wang, S. et al. Operon prediction by decision tree classifier based on vprsm. In 2009 3rd International Conference on Bioinformatics and Biomedical Engineering, 1–4 (IEEE, 2009).

[18] Parks, D. H. et al. Recovery of nearly 8, 000 metagenome-assembled genomes substantially expands the tree of life. Nature Microbiology 2, 1533–1542 (2017). URL 10.1038/s41564-017-0012-7.

[19] Assaf, R., Xia, F. & Stevens, R. Detecting operons in bacterial genomes via visual representation learning. Scientific Reports 11, 2124 (2021).

[20] Tomar, T. S., Dasgupta, P. & Kanaujia, S. P. Operon finder: a deep learning-based web server for accurate prediction of prokaryotic operons. Journal of Molecular Biology 435, 167921 (2023).

[21] Wattam, A. R. et al. Improvements to patric, the all-bacterial bioinformatics database and analysis resource center. Nucleic acids research 45, D535–D542 (2017).

[22] Mering, C. v. et al. String: a database of predicted functional associations between proteins. Nucleic Acids Res. 31, 258– 261 (2003).

[23] Szklarczyk, D. et al. The string database in 2021: customizable protein–protein networks, and functional characterization of user-uploaded gene/measurement sets. Nucleic acids research 49, D605–D612 (2021).

[24] Zaidi, S. S. A., Kayani, M. U. R., Zhang, X., Ouyang, Y. & Shamsi, I. H. Prediction and analysis of metagenomic operons via metaron: a pipeline for prediction of metagenome and whole-genome operons. BMC genomics 22, 1–14 (2021).

[25] Hyatt, D. et al. Prodigal: prokaryotic gene recognition and translation initiation site identification. BMC bioinformatics 11, 1–11 (2010).

[26] Database resources of the national center for biotechnology information. Nucleic acids research 46, D8–D13 (2018).

[27] Tamames, J., Casari, G., Ouzounis, C. & Valencia, A. Conserved clusters of functionally related genes in two bacterial genomes. Journal of molecular evolution 44, 66–73 (1997).

[28] Kanehisa, M., Furumichi, M., Tanabe, M., Sato, Y. & Morishima, K. Kegg: new perspectives on genomes, pathways, diseases and drugs. Nucleic acids research 45, D353–D361 (2017).

[29] Steinegger, M. & Söding, J. MMseqs2 enables sensitive protein sequence searching for the analysis of massive data sets. Nature Biotechnol. 35, 1026–1028 (2017).

[30] Kiesel, A. et al. The bamm web server for de-novo motif discovery and regulatory sequence analysis. Nucleic acids research 46, W215–W220 (2018).

[31] Ge, W., Meier, M., Roth, C. & Söding, J. Bayesian markov models improve the prediction of binding motifs beyond first order. NAR Genomics and Bioinformatics 3, lqab026 (2021).

[32] Finn, R. D. et al. The Pfam protein families database: towards a more sustainable future. Nucleic Acids Res. 44, D279–D285 (2016).

[33] Schmalz, X., Biurrun Manresa, J. & Zhang, L. What is a bayes factor? Psychological methods 28, 705 (2023).

[34] Salgado, H., Moreno-Hagelsieb, G., Smith, T. F. & Collado-Vides, J. Operons in escherichia coli: genomic analyses and predictions. Proceedings of the National Academy of Sciences 97, 6652–6657 (2000).

[35] Chuang, L.-Y., Chang, H.-W.Tsai, J.-H. & Yang, C.-H. Features for computational operon prediction in prokaryotes. Briefings in functional genomics 11, 291–299 (2012).

[36] Griffiths, A. J. An introduction to genetic analysis (Macmillan, 2005).

[37] Mushegian, A. R. & Koonin, E. V. Gene order is not conserved in bacterial evolution. Trends in genetics: TIG 12, 289–290 (1996).

[38] Zhang, R. Spacedust: de novo discovery of conserved gene clusters from large-scale bacterial genome sets. https://github.com/soedinglab/spacedust (2023).

[39] Saito, T. & Rehmsmeier, M. The precision-recall plot is more informative than the roc plot when evaluating binary classifiers on imbalanced datasets. PloS one 10, e0118432 (2015).

[40] Stein, L. Generic feature format version 3 (gff3). Seq. Ontol. Proj 1 (2013).

[41] Tierrafría, V. H. et al. Regulondb 11.0: Comprehensive high-throughput datasets on transcriptional regulation in escherichia coli k-12. Microbial Genomics 8, 000833 (2022).

[42] Pedreira, T., Elfmann, C. & Stülke, J. The current state of subti wiki, the database for the model organism bacillus subtilis. Nucleic Acids Research 50, D875–D882 (2022).

[43] Sierro, N., Makita, Y., de Hoon, M. & Nakai, K. Dbtbs: a database of transcriptional regulation in bacillus subtilis containing upstream intergenic conservation information. Nucleic acids research 36, D93–D96 (2008).

[44] Chetal, K. & Janga, S. C. Operomedb: a database of condition-specific transcription units in prokaryotic genomes. BioMed research international 2015 (2015).

