## Supplementary Materials for "UniOP: a universal operon prediction for high-throughput prokaryotic (meta-)genomic data using intergenic distance"

### Self-supervised training to predict operons in complete genomes

**Selection of reference database.** As our reference, we utilized a comprehensive dataset of 4683 representative prokaryotic genomes from the RefSeq database, all at chromosome or complete assembly level. The conservation of a gene pair in distantly related genomes typically provides more significant insights into the importance and evolutionary stability of that gene pair than its conservation in closely related genomes. To eliminate redundancy and enhance the efficiency of detecting conserved gene clusters in the reference genomes, we implemented a systematic procedure that accounts for the evolutionary distances between the query genome and the reference genomes.

The following steps outline this process in detail. Initially, we calculate pairwise evolutionary distances using the mash program [1] with default parameters, enabling comprehensive comparisons between the query genome and the entire reference database. By analyzing these evolutionary distances, we select genomes with distances ranging between 0 and 1 (without 0 and 1). This range ensures that the chosen reference genomes exhibit sufficient evolutionary relatedness to the query genome while excluding highly dissimilar genomes. Next, we adopt a selection strategy to minimize redundancy that selects a maximum of two genomes at each unique distance value identified in the previous step. This restriction prevents the overrepresentation of genomes that share a close evolutionary distance with the query genome in the final reference databases. By imposing this limit, we encourage diversity and minimize potential bias.

Through this process, we effectively curate a customized reference database for each query genome, optimizing the accuracy and relevance of subsequent analyses. This approach ensures that each query genome is compared against a diverse and representative set of reference genomes, enhancing the reliability of our findings on conserved gene pairs.

**Search for matching clusters of genes between $g$ and all selected reference genomes $\mathcal{G}$.** For a query genome $g$, suppose we have a selected reference set $\mathcal{G}$ with $G$ genomes. We first employ Spacedust to systematically identify gene clusters in $g$ across $\mathcal{G}$. Next, we construct a raw conservation matrix $Y_{gij}$ with entries $y_{gij,g'}$, where each row represents a reference genome $g'$, and each column represents a same-strand neighboring gene pair $i$ and $j = i + 1$ in genome $g$. Set $y_{gij,g'} = 1$ if proteins $i$ and $j$ are both contained in a cluster matched to a cluster in genome $g'$. Otherwise, set $y_{gij,g'} = 0$. We center the explanatory variables,

$$y_{gij,g'} \leftarrow y_{gij,g'} - \frac{1}{N_g} \sum_{i'=1}^{N_g} y_{gi'j,g'} \,, \tag{1}$$

but *we do not normalize them*. In this way, for closely related genomes with 10% of disrupted neighboring gene pairs, $y_{gi'j,g'}$ will be 0.1 for the uninterrupted pairs and -0.9 for the interrupted ones. Conversely, for distantly related genomes with 10% of conserved neighboring gene pairs, $y_{gi'j,g'}$ will be 0.9 for the conserved pairs and -0.1 for the other 90% of pairs. Here, $N_g$ is the total number of same-strand genes in genome $g$.

Additionally, we create a refined conservation matrix $Y'_{gij}$ by substituting $y_{gij,g'}$ values with the UniOP predictions for the gene pair $i'$ and $j' = i'+1$ in reference genome $g'$. If $y_{gij,g'} = 1$, then the value is replaced by the UniOP prediction for the gene pair ($i'$,$j'$). Otherwise, the values remain the same.

**Assign pseudo labels using UniOP.** We denote by $z_{gij} = 1$ the case that two genes $i$ and $j = i+1$ from the same strand of genome $g$ are part of the same operon, and with $z_{gij} = 0$ that they are not. The $z_{gij}$ are the labels we would like to predict. We estimate the probability of two neighboring genes being part of the same operon using the distance model, UniOP. Subsequently, we assign *pseudo labels* to these genes based on their probabilities. For a fraction $\alpha q$ of genes with highest probability $p(z_{gij}=1|d)$, we set $z_{gij} = 1$. Conversely, we set $z_{gij} = 0$ for a fraction $\alpha(1-q)$ of genes with lowest probability $p(z_{gij} = 1|d)$. The value of $\alpha$ determines the proportion of gene pairs that will receive a label, so a smaller $\alpha$ results in fewer but more reliable pseudo labels. In this case, we set $\alpha = 0.5$. We use these gene pairs with highly confident labels to train two logistic regression models based on $Y_{gij}$ and $Y'_{gij}$, respectively.

**Logistic regression prediction of the probability for genes being the same operon.** With the logistic function $\sigma(x) = 1/(1+e^{-x})$, the probability for $z_{gij}=1$ under the logistic regression model is

$$p(z_{gij}=1|\mathbf{y}_{gij}, \mathbf{w}_g) = \sigma(\mathbf{w}_g^{\mathrm{T}}\mathbf{y}_{gij} + \delta_g)\,, \tag{2}$$

where $\mathbf{y}_{gij}$ is the vector of dimension $G$ with entries $y_{gij,g'}$, $G$ is the number of selected reference genome set $\mathcal{G}$, $g'$ is all genomes in $\mathcal{G}$. Therefore,

$$p(z_{gij}|\mathbf{y}_{gij}, \mathbf{w}_g) = \sigma(\mathbf{w}_g^{\mathrm{T}}\mathbf{y}_{gij} + \delta_g)^{z_{gij}} \left(1 - \sigma(\mathbf{w}_g^{\mathrm{T}}\mathbf{y}_{gij} + \delta_g)\right)^{1-z_{gij}}\,. \tag{3}$$

To find the model parameters, we need to maximize the regularized log-likelihood,

$$\mathsf{LL}_{\mathsf{reg}}(\mathbf{w}_g) = \sum_{i=1}^{N_g-1} \left(z_{gij}\ln\sigma(\mathbf{w}_g^{\mathrm{T}}\mathbf{y}_{gij} + \delta_g) + (1-z_{gij})\ln\left(1 - \sigma(\mathbf{w}_g^{\mathrm{T}}\mathbf{y}_{gij} + \delta_g)\right)\right) - R(\mathbf{w}_g) \tag{4}$$

where $R(\mathbf{w}_g)$ is a regularizer to limit the model's complexity and include prior knowledge. A simple regularizer is $R(\mathbf{w}_g) = \lambda\,\|\mathbf{w}_g\|^2$. Importantly, since $y_{gij,g'} < 0$ is always evidence against $z_{gij,g'} = 1$ and $y_{gij,g'} > 0$ is evidence for it, regression coefficients cannot be negative,

$$w_{g,g'} \geq 0\,. \tag{5}$$

We need to learn $G$ parameters, $\mathbf{w}_g = (w_{g,g'} : g' \in \mathcal{G})$, one per genome, and bias term $\delta_g$.

**Combined model using distance and conservation.** Once the regression model is trained, we want to include the intergenic distance information. We can do that by making the bias $\delta_g$ depend on the intergenic distance, $\delta_g(d_{gij})$. Imagine that all weights were zero, then we would want to predict the probability $p(z_{gij}=1|\mathbf{y}_{gij}, \mathbf{w}_g=\mathbf{0}, d_{gij}) = p(z_{gij}=1|d_{gij})$, or equivalently

$$\delta_g(d_{gij}) = \ln\frac{p(z_{gij}=1|\mathbf{y}_{gij}, \mathbf{w}_g=\mathbf{0}, d_{gij})}{p(z_{gij}=0|\mathbf{y}_{gij}, \mathbf{w}_g=\mathbf{0}, d_{gij})} = \ln\frac{p(z_{gij}=1|d_{gij})}{p(z_{gij}=0|d_{gij})}\,. \tag{6}$$

This fixes the bias $\delta_g(d_{gij})$ as a function of $d_{gij}$. To allow the regression model to deviate slightly from the prior probabilities given by the distance-based model, we add a global bias $\delta_g$ to the distance-dependent one. We therefore compute the probabilities for $z_{gij} = 1$ given the search results $\mathbf{y}_{gij}$ *and the intergenic distance* $d_{gij}$, as

$$p(z_{gij}=1|\mathbf{y}_{gij}, \mathbf{w}_g, d_{gij},) = \sigma\left(\mathbf{w}_g^{\mathrm{T}}\mathbf{y}_{gij} + \ln\frac{p(z_{gij}=1|d_{gij})}{p(z_{gij}=0|d_{gij})}\right)\,. \tag{7}$$

with the fixed weights from the regression without distance information.

**Hyperparameters optimization of the conservation model** We utilized L2-regularization to determine the optimal hyperparameters for the LR model. These hyperparameters, which include the $\lambda$, learning rate $lr$, and the number of iterations $n$, were tuned on the *E.coli* dataset using 5-fold cross-validation. To accomplish this, a grid search was performed with $\lambda$ values in the range [0.1, 1, 10, 100], $lr$ values in [0.001, 0.01, 0.1, 1] and $n$ values in [100, 150, 200, 300]. GridSearchCV function from sklearn 1.2.2 was employed for this purpose. The best combination of hyperparameters, which yield the optimal prediction of distance-based pseudo labels by the regression model, was selected. The final values for $\lambda$, $lr$, and $n$ were determined as 10, 0.01 and 100, respectively. These values were applied to other genomes without further optimization to ensure generality.

**Table 1. Comparison between annotated and estimated $q$ values across the ten genomes.** The ratio is defined as $\log(q_{\text{est}}/q_{\text{annot}})$.

| species | $q_{\text{annot}}$ | $q_{\text{est}}$ | ratio | category |
|---|---|---|---|---|
| *Escherichia coli* | 0.58 | 0.574 | -0.011 | Group 1 |
| *Mycobacterium tuberculosis* | 0.506 | 0.485 | -0.043 | Group 1 |
| *Bacillus subtilis* | 0.596 | 0.628 | 0.053 | Group 1 |
| *Campylobacter jejuni* | 0.686 | 0.748 | 0.087 | Group 1 |
| *Synechococcus elongatus* | 0.41 | 0.452 | 0.097 | Group 1 |
| *Salmonella enterica* | 0.497 | 0.586 | 0.164 | Group 2 |
| *Sinorhizobium meliloti* | 0.42 | 0.531 | 0.236 | Group 2 |
| *Eggerthella lenta* | 0.469 | 0.631 | 0.298 | Group 2 |
| *Clostridioides difficile* | 0.414 | 0.78 | 0.643 | Group 3 |
| *Clostridium beijerinckii* | 0.286 | 0.722 | 0.926 | Group 3 |

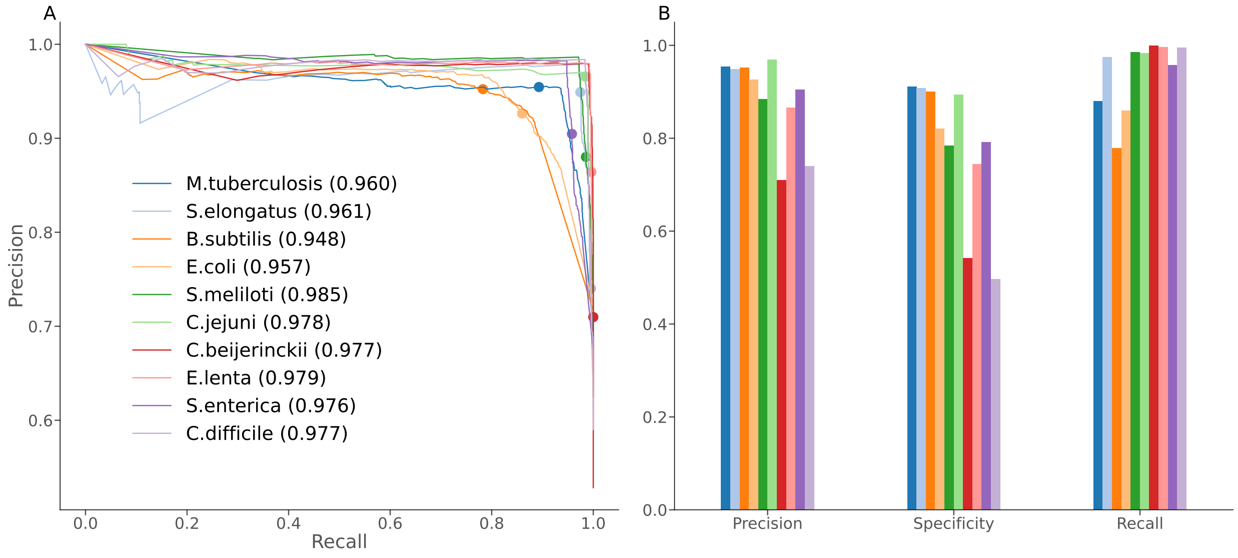


**Fig. 1. Evaluation performance of UniOP over ten genomes. A**. Precision-Recall curves. Each dot on the curve indicates the precision and recall values at the threshold of 0.5, with the corresponding precision and recall bar plotted in **B**.

**Table 2. Human gut MAGs.** #operons refers to the UniOP prediction, with the total number of operons across all phyla summing to 1,603,492. %annotated pairs indicates the proportion of gene pairs within each phylum annotated by the same KEGG module(s). %negatives represents the proportion of gene pairs within each phylum annotated by the different KEGG module(s). %operonic genes is defined as the proportion of genes covered by predicted operons relative to the total number of genes across all MAGs within each phylum.

| Phylum | #MAGs | #adjacent pairs | #positives | #negatives | %annotated pairs | #operons | %operonic genes |
|---|---|---|---|---|---|---|---|
| *Firmicutes_A* | 1229 | 2,346,794 | 210,610 | 64,187 | 0.117 | 625,877 | 0.786 |
| *Actinobacteriota* | 599 | 789,703 | 91,334 | 25,507 | 0.148 | 235,719 | 0.686 |
| *Bacteroidota* | 436 | 836,464 | 47,955 | 20,230 | 0.082 | 255,537 | 0.735 |
| *Firmicutes* | 420 | 735,890 | 68,749 | 17,248 | 0.117 | 183,781 | 0.754 |
| *Proteobacteria* | 280 | 621,493 | 73,923 | 20,980 | 0.153 | 177,730 | 0.595 |
| *Firmicutes_C* | 114 | 172,317 | 19,069 | 6990 | 0.151 | 37,960 | 0.846 |
| *Campylobacterota* | 43 | 60,321 | 5749 | 3322 | 0.150 | 16,396 | 0.807 |
| *Verrucomicrobiota* | 39 | 69,147 | 4334 | 2747 | 0.102 | 20,700 | 0.701 |
| *Fusobacteriota* | 32 | 54,603 | 5391 | 1541 | 0.127 | 14,127 | 0.830 |
| *Desulfobacterota* | 25 | 44,489 | 4371 | 1744 | 0.137 | 13,740 | 0.695 |
| *Spirochaetota* | 18 | 27,891 | 2610 | 904 | 0.126 | 8906 | 0.674 |
| *Cyanobacteria* | 11 | 13,266 | 984 | 509 | 0.113 | 4124 | 0.563 |
| *Thermoplasmatota* | 11 | 12,639 | 1787 | 418 | 0.174 | 3888 | 0.557 |
| *Methanobacteriota* | 10 | 11,970 | 1259 | 516 | 0.148 | 3860 | 0.656 |
| *Halobacteriota* | 2 | 4438 | 440 | 96 | 0.121 | 1147 | 0.418 |

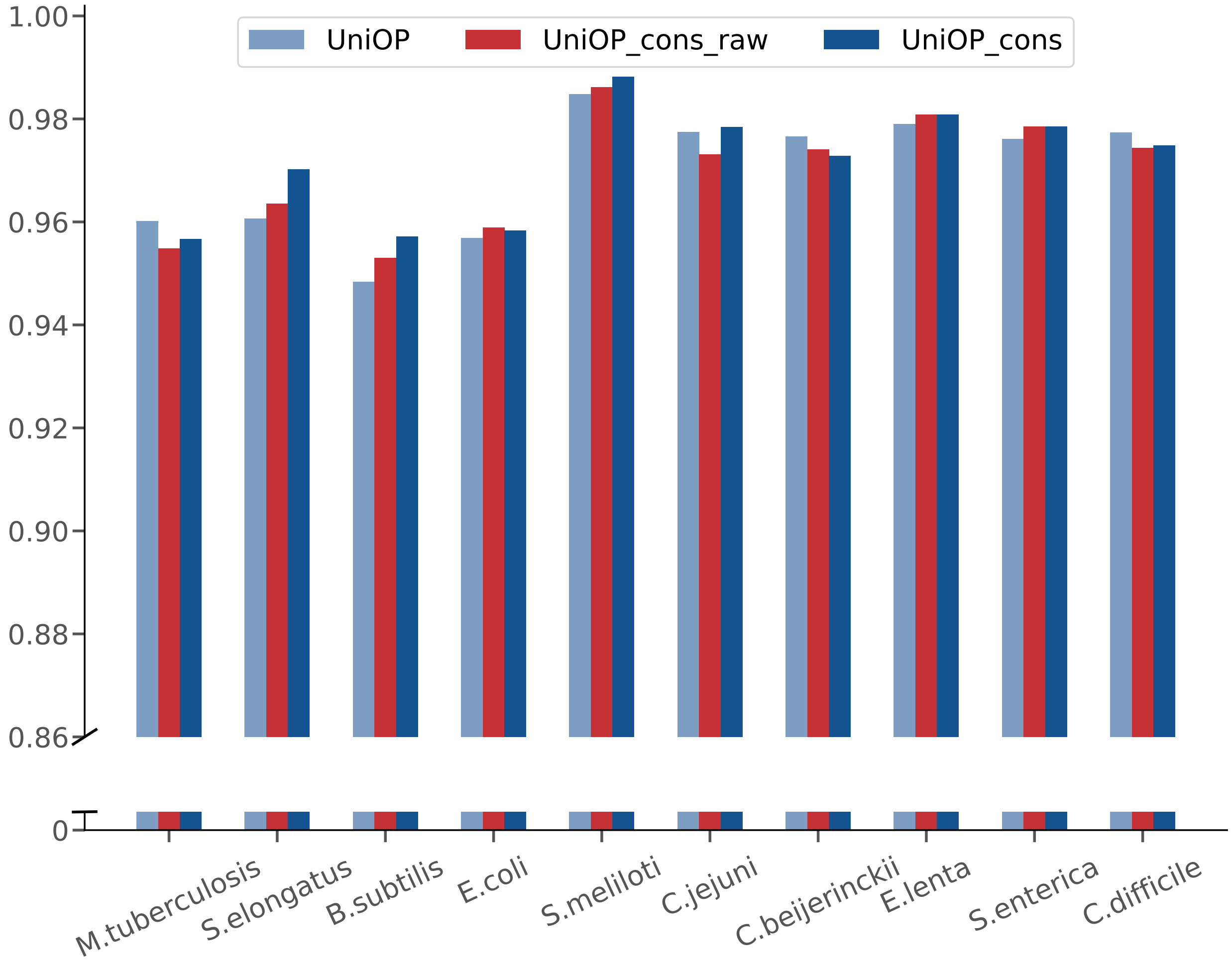


**Fig. 2. Predictive performance of UniOP by combining it with two different conservation matrices.** UniOP_cons_raw (red bar) represents a combination of the predictions from UniOP and those from LR trained on raw conservation matrix. In contrast, UniOP_cons (dark blue bar) represents a combination of the predictions from UniOP and those from LR trained on the refined conservation matrix.

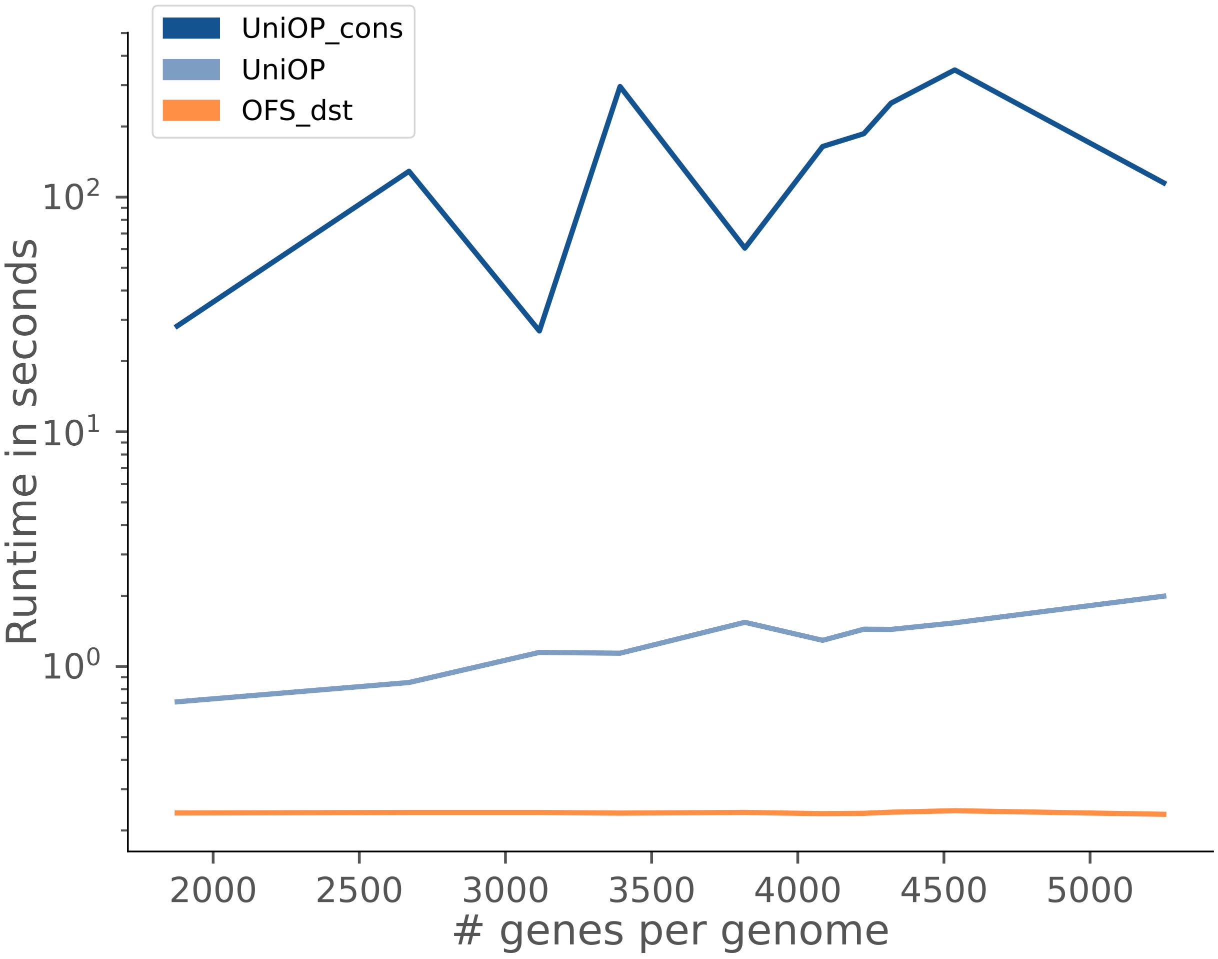


**Fig. 3. Comparison of runtime in the function of the number of genes per genome for UniOP, UniOP_cons (predicting operons by incorporating conserved gene cluster information into UniOP), and OFS_dst over ten genomes.** The y-axis is the runtime in seconds. The x-axis is the number of genes per genome.

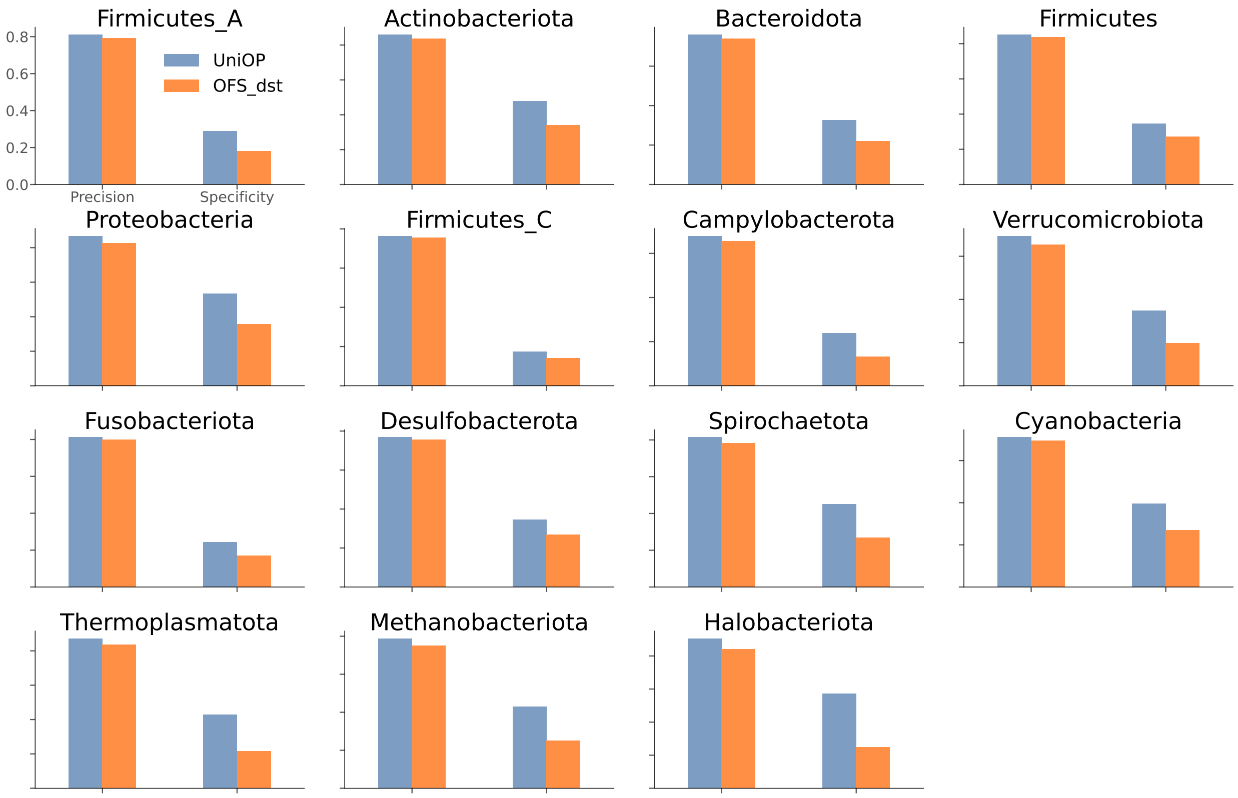


**Fig. 4. Precision and Specificity pf UniOP and OFS_dst for MAGs.**

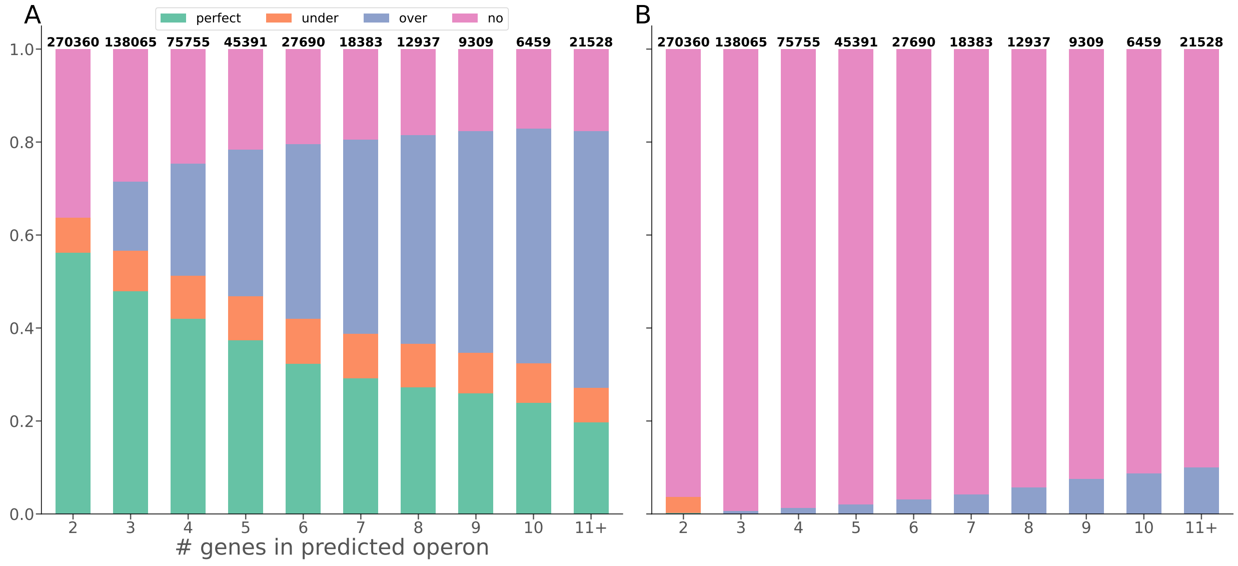


**Fig. 5. Conservation results for MAGs. A** The proportion of four operon conservation classes in original MAGs. **B** The proportion of four operon conservation classes in shuffled MAGs.

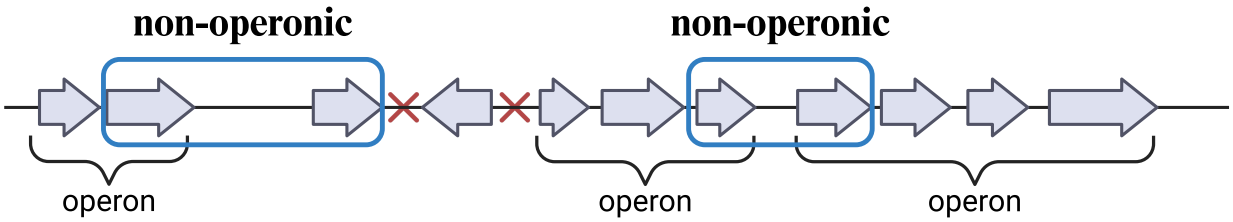


**Fig. 6. Labeling of non-operonic gene pairs.** Gene pairs are classified as non-operonic pairs if the first and last genes of operons are on the same strand with their upstream and downstream genes, respectively.

### References

[1] Ondov, B. D. *et al.* Mash: fast genome and metagenome distance estimation using minhash. *Genome biology* **17**, 1–14 (2016).
